# Towards an Unconscious Neurotherapy for Common Fears

**DOI:** 10.1101/170183

**Authors:** Vincent Taschereau-Dumouchel, Aurelio Cortese, Toshinori Chiba, J.D. Knotts, Mitsuo Kawato, Hakwan Lau

## Abstract

Can evolutionarily ‘hardwired’ fear responses, e.g. for spiders and snakes, be reprogramed unconsciously in the human brain? Currently, exposure therapy is amongst the most effective treatments for anxiety disorders^1^, but this intervention is subjectively aversive to patients, and rates of premature attrition from treatment have been reported to be as high as 70%^2^. Here we introduce a novel method to bypass the subjective unpleasantness in conscious exposure, by directly pairing monetary reward with unconscious occurrences of decoded representations of naturally feared objects in the brain. The typical way to identify multivoxel functional magnetic resonance imaging (fMRI) representations for feared objects involves repeated presentations of the relevant images explicitly to subjects. However, for our potential treatment method to be effective in actual clinical settings, we need to decode fear representations *without* triggering excessively aversive reactions which may cause patients to dropout from treatments prematurely. Here we overcome this challenge by capitalizing on recent advancements in fMRI decoding techniques: We employed a method called hyperalignment^3,4^ to infer the relevant representations of feared objects for a designated participant based on data from other ‘surrogate’ participants. This way the procedure completely bypasses the need for the conscious encountering of feared objects. We demonstrate that our method can lead to reliable reductions in physiological fear responses measured by skin conductance as well as amygdala hemodynamic activity. Not only do these results raise the intriguing possibility that naturally occurring fear can be ‘re-programmed’ outside of conscious awareness, importantly they also created the rare opportunity for a psychological intervention of this nature to be tested rigorously in a double-blind placebo-controlled fashion. This may pave the way for a novel treatment method, combining the appealing rationale and proven efficacy of conventional psychotherapy with the rigor and leverage of clinical neuroscience.

---

Recent advancements in neuroimaging and machine learning have made it possible for us to identify specific representations of commonly and naturally feared objects in in the human brain^5^. Here we tested the hypothesis that despite the supposed deep evolutionary basis of these neural representations^6^, we can reprogram their associations to reduce the relevant fear responses. Previously, using closed-loop decoded fMRI neural-reinforcement, we have shown that physiological fear responses can be reduced by pairing rewards with the occurrences of decoded object representations^7^. However, in that study, the artificial objects were feared only because they have been experimentally conditioned with electric shocks, and that procedure was itself conscious. Here, we tested if our neural-reinforcement method may apply to naturally occurring fear in everyday stimuli, e.g. images of spiders or snakes, completely without participants’ awareness. This also has the advantage of mimicking more closely the actual clinical context in which direct visual presentations of feared objects may not be easily tolerated by patients.

In order to construct across-subjects machine learning decoders for some of the most commonly feared animals, we designed an experiment (Day 0: Decoder Construction session) in which normal healthy participants viewed 3600 images of 30 different animals and 10 inanimate objects (Fig. 1a). The multivoxel fMRI responses to these individual images were recorded in the fMRI scanner and extracted at the single-trial level, using conventional analytic procedures (see Methods).

**Figure 1.**
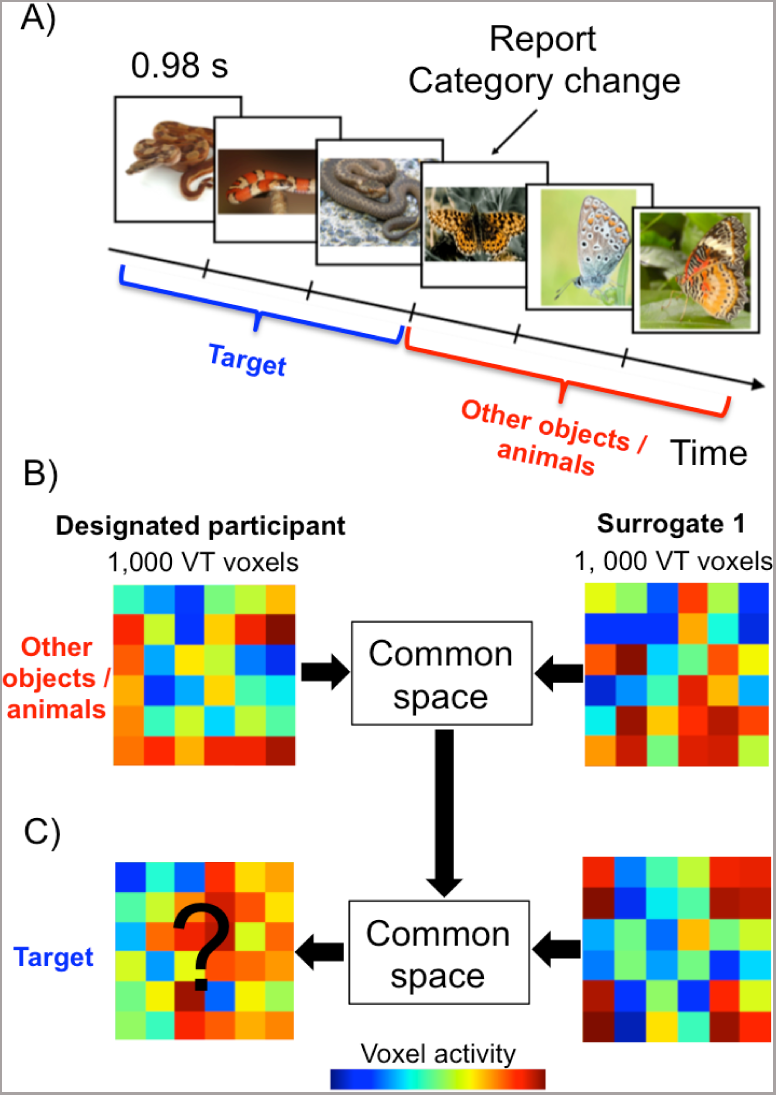
Hyperalignment Decoder Construction. A) An example sequence of events in the Hyperalignment Decoder Construction session. B) To mimic the situation wherein patients are not exposed to Target images (to avoid excessive fear), the construction of Hyperlignment decoders was based on the data from ‘surrogate’ participants. To do so, we ‘hyperaligned’ voxels in the ventral temporal (VT) area between a designated participant and surrogates into a ‘common space’, using the representations for the 39 Non-target categories. C) Through this space, we created a Hyperalignment Decoder for a Target category in the designated participant based on the surrogates’ representations for that Target category.

We then capitalized on a novel method of across-subjects multivoxel analysis known as hyperalignment, which allowed us to compare and translate patterns of fMRI activity between participants^3,4^. Using this method, we exploited the data from as many as 29 ‘surrogate’ participants (who viewed the Target feared category) to construct the Hyperalignment Decoder for a designated participant as if they were never exposed to the Target images (as would be ideal in a clinical setting) (Fig. 1b). To do so, an abstract common space was derived from the voxel activity in the ventral temporal cortex (fusiform, inferior temporal, lingual/parahippocampal cortex)^3^ of all participants based on all image categories except for the Target category (e.g., snake). This allowed us to infer the decoders for the designated participant based on the decoders of ‘surrogate’ participants (extended data Fig. 1). Specifically, we trained a decoder to discriminate multivoxel patterns for the Target category from patterns for all of the other Non-target categories in the ‘surrogate’ participants (Target vs Non-target), and through transformations via the common space we inferred what such a decoder would be for the designated participant. We call this the Hyperalignment Decoder.

One may worry that such an indirect inference strategy may only provide limited efficiency. However, for each designated participant, this procedure can benefit from the data of as many as 29 ‘surrogate’ participants. As such we harness the power of a much larger amount of data to train the decoders than in conventional (i.e., within-subject) fMRI decoding. As in previous reports^3^, these Hyperalignment Decoders displayed decoding accuracies (M = 82.4 %; S.E.M. = 0.27%) even higher (*t*(39) = 9.55; *P* < .0001; Wilcoxon signed-rank test: z = 4.76, p < .0001) than accuracies obtained with traditional within-subject decoders trained by presenting images to the participants themselves (M = 71.7%; S.E.M. = 1.01%) (Fig. 2b). These results are also in accordance with previous data indicating that the topography of voxels’ selectivity appears to be preserved by hyperalignment^3^ (Fig. 2a and extended data Table 1).

**Figure 2.**
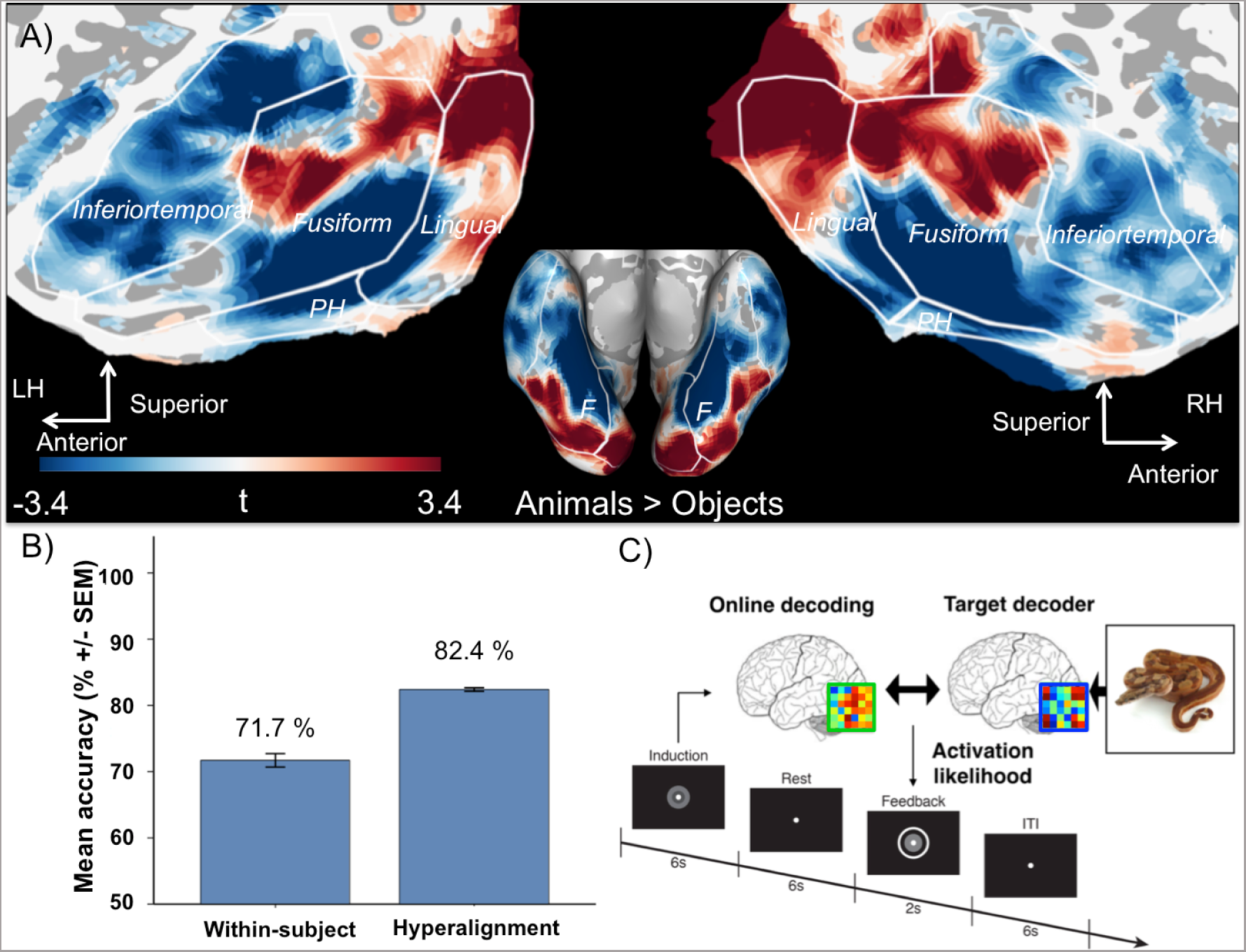
Classification accuracies and Neural-Reinforcement procedure. A) Voxels contributing to the Hyperalignment Decoders: plotted are the t-values of the voxels’ weights, which are consistent with a lateral-to-medial animacy continuum^3,8^ (critical *t*-value = 3.4 for p < .001, uncorrected). B) Hyperalignment Decoders built using 29 ‘surrogate’ participants reached higher accuracies than conventional within-subject methods. C) During Neural-Reinforcement, online decoding was used to reinforce occurrences of the Target (but not Control) multivoxel representation. The feedback was proportional to the likelihood of the Target being represented, and to monetary gain. Crucially, participants and experimenters were both unaware of the identity of the Target category throughout the experiment.

**Extended Data Table 1.** Spatial distribution of the sparse-logistic-regression weights.

| **Contrast** | **Label** | **$P_{\text{FDR}}$**<br>**cluster-corrected** | **Peak**<br>**$t$-score** | **Peak voxel MNI**<br>**coordinates** |
| --- | --- | --- | --- | --- |
| **Object >**<br>**Animal** | Left fusiform | <.001 | 15.69 | [-26,-50,-12] |
|  | Right fusiform | <.001 | 12.08 | [30,-38,-16] |
|  | Left inferior<br>temporal | <.001 | 6.75 | [-52,-62,-12] |
|  | Left anterior<br>temporal | .006 | 5.25 | [-44,-12,-40] |
|  | Right inferior<br>temporal | .044 | 4.47 | [60,-22,-34] |
| **Animal >**<br>**Object** | Left and right<br>lingual | <.001 | 10.62 | [-6,-92,-12] |
|  | Right fusiform | .004 | 6.85 | [44,-50,-22] |
|  | Left fusiform | .009 | 5.81 | [-44,-56,-22] |

Importantly, whether the designated participant presented a similar fear profile (over all 40 categories of animals/objects) to the average fear profile of all participants in the hyperalignment did not modulate the accuracy of the Hyperalignment Decoder (Extended Data Fig. 5). This suggests that our method may be promising even for patients with atypical fear profiles. Preliminary data with actual patients diagnosed with specific phobias further support this claim (Extended Data Fig. 6).

We then conducted five days of our Neural-Reinforcement procedure (Fig. 3a) using these Hyperalignment Decoders in 17 participants who presented high (subclinical) levels of fear for at least two animal categories in our database (see Methods for details on the selection process). For each participant, one of the feared animal categories was selected randomly by the computer to be the Target of the intervention, while the other of the feared animal categories acted as a within-subject control to allow us to determine the specificity of the intervention. Physiological fear response was assessed before and after Neural-Reinforcement sessions by presenting images of the two feared categories (Target & Control) and images of a neutral category (Baseline condition) while measuring skin conductance response and amygdala hemodynamic response.

**Figure 3.**
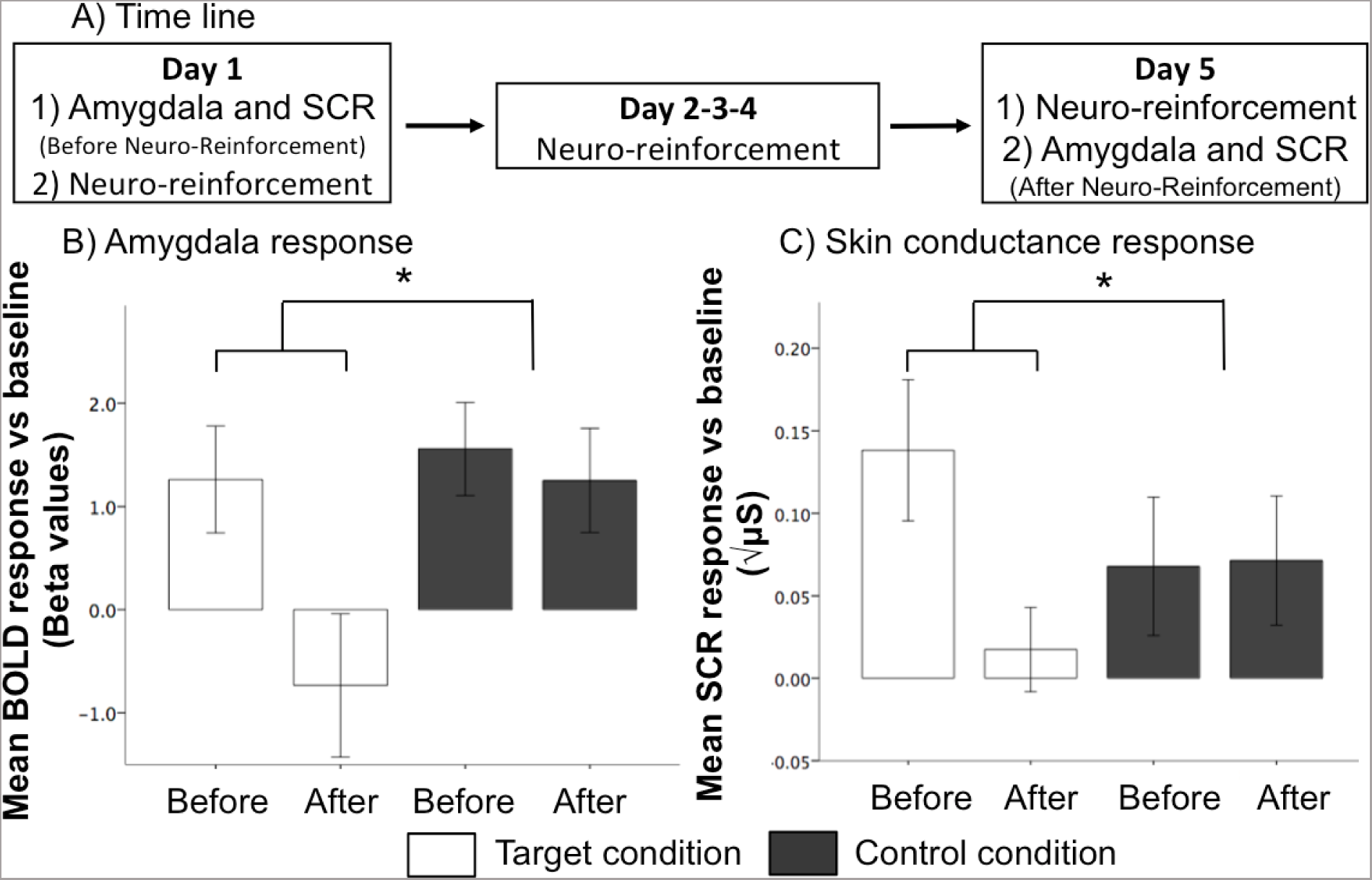
Decrease in physiological fear responses following Neural-Reinforcement. A) To assess changes in physiological fear responses, on Days 1 and 5, participants viewed images of two animal categories that they feared (Target and Control) and one animal of which they were not afraid (Baseline). B) The results indicate a significant decrease of amygdala response for the Target condition while the Control condition remained unchanged by the procedure. C) Likewise, the results indicate a significant decrease in skin conductance response in the Target condition and no decrease in the Control condition. BOLD: Blood-oxygen-level dependant. SCR: Skin conductance response. Error bars are +/- 1 S.E.M. \**P* < .05.

In each trial during the Neural-Reinforcement procedure (Fig. 2c), the Hyperalignment Decoder was applied to fMRI images in real time to determine the likelihood that the pattern of activity corresponding to the Target category was represented in the brain (see Extended Data Fig. 2). This information was visually fed back to participants via varying the size of a disc image from trial to trial, which was directly proportional to the amount of money the subject would earn on a trial. Following previous procedures^7,9–12^, participants were explicitly told about the association between disc size and monetary reward, but received no instructions as to what brain activity patterns were necessary to maximize the size of the disc.^7,9–12^. Despite this, participants were able to learn to activate the Target representation with statistical consistency above chance (see Extended Data Fig. 2a and Supplementary Discussion). This process was thus conducted in a double-blinded fashion, as neither the participants nor the experimenters were aware of the identity of the Target category.

Confirming our hypothesis, we found a specific reduction of physiological fear response for the Target category after Neural-Reinforcement (Fig. 3): Amygdala response for the Target condition decreased (*t*(16) = 2.41; *P* = .028), but remained unchanged for the Control condition (*t*(16) = .40; *P* = .69, and showed a significant time X condition interaction *F*(1,16) = 5.57; *P* = .031). Likewise, we observed a significant decrease in skin conductance response for the Target condition (*t*(122) = −2.48; *P* = .014), but not the Control condition (*t*(122) = .016; *P* = .99, significant time X condition interaction: *F*(1,244) = 2.13; *P* = .033).

Importantly, at the end of the procedure, participants were unable to guess the identity of the Target category (47% accuracy in a two-alternative forced-choice question), and reported strategies for maximizing rewards generally unrelated to the Target and purpose of the procedure (Extended data Table 2 for induction strategies reported by participants). This confirms that our treatment effects can be obtained outside of participants’ conscious awareness (see also Extended Data Fig. 3).

**Extended Data Table 2.**
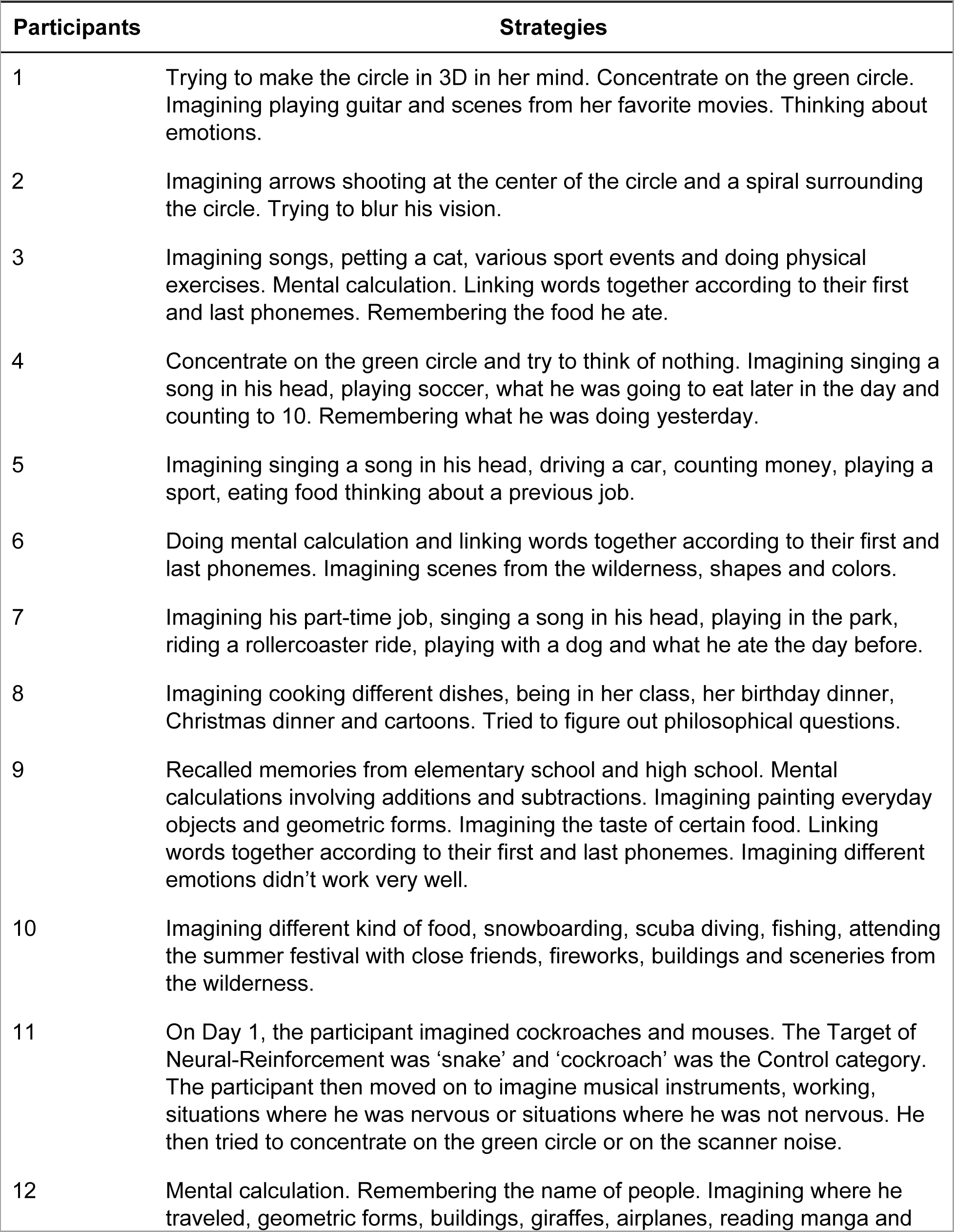
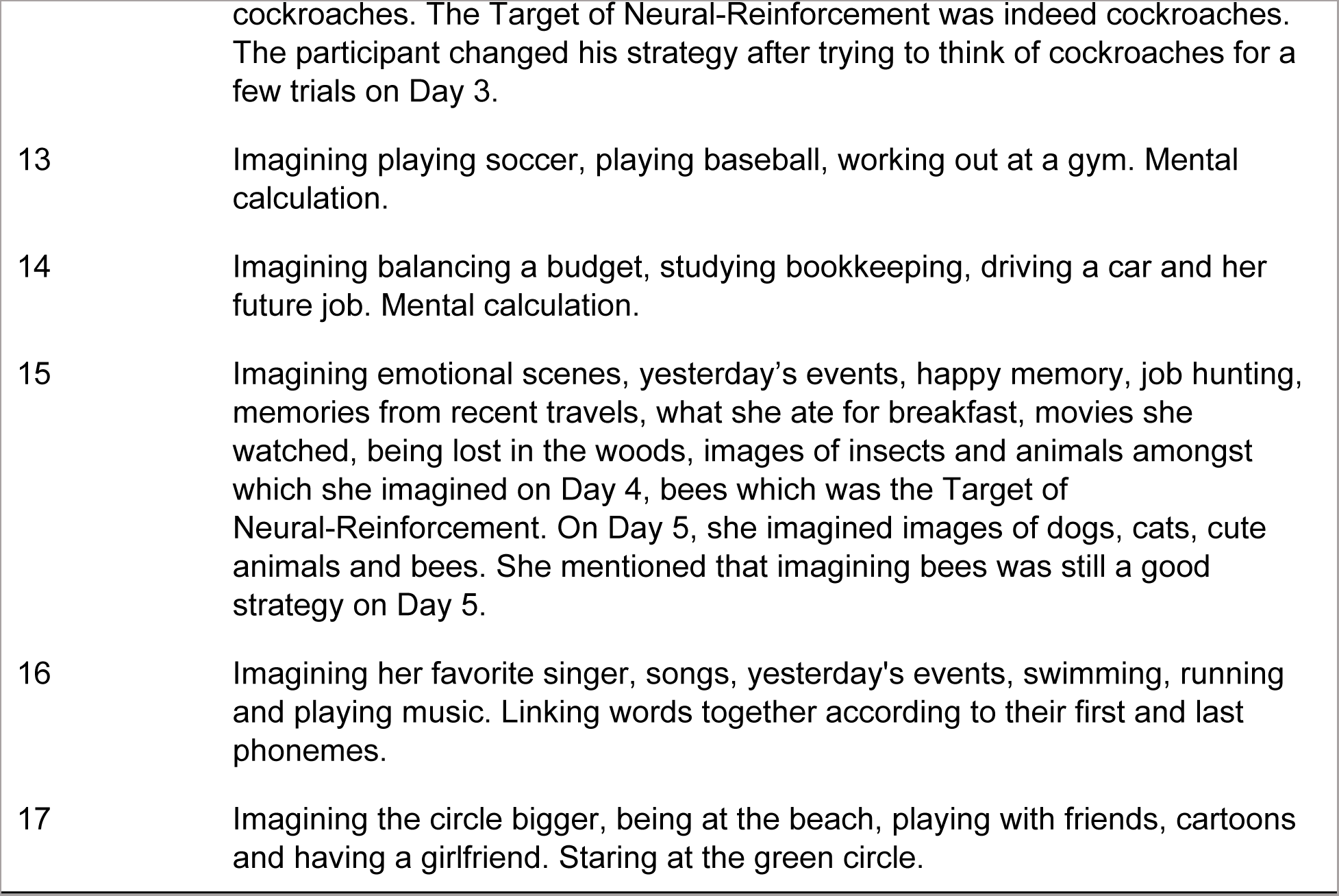
Strategies reported by participants.

What could be the mechanisms underlying these results? To better understand the nature of Neural-Reinforcement, we conducted information transmission analyses^7,9,10^ to investigate if multivoxel patterns in other brain regions can predict, on a trial-by-trial basis, the likelihood of Target representation in the ventral temporal cortex. These analyses indicated that, during Neural-Reinforcement, the voxels tracking the likelihood of Target representation were primarily contained in the fusiform, inferior temporal, and lingual cortex (Fig. 4). These results were compared to normal conscious viewing of Target images (Day 0: Decoder Construction session), wherein the voxels tracking the decoders’ likelihood were distributed more broadly in the fusiform, and lingual regions, as well as outside of the ventral temporal cortex in areas such as the amygdala (Extended Data Fig. 4), cuneus, and parietal and occipital regions (Fig. 4). Overall, these observations are consistent with our previous report that, during Neural-Reinforcement, the induced Target representations were relatively localized and disconnected from the rest of the fear-related circuitry. Such a disconnect may be an important aspect of our fear reduction procedure^7^.

**Figure 4.**
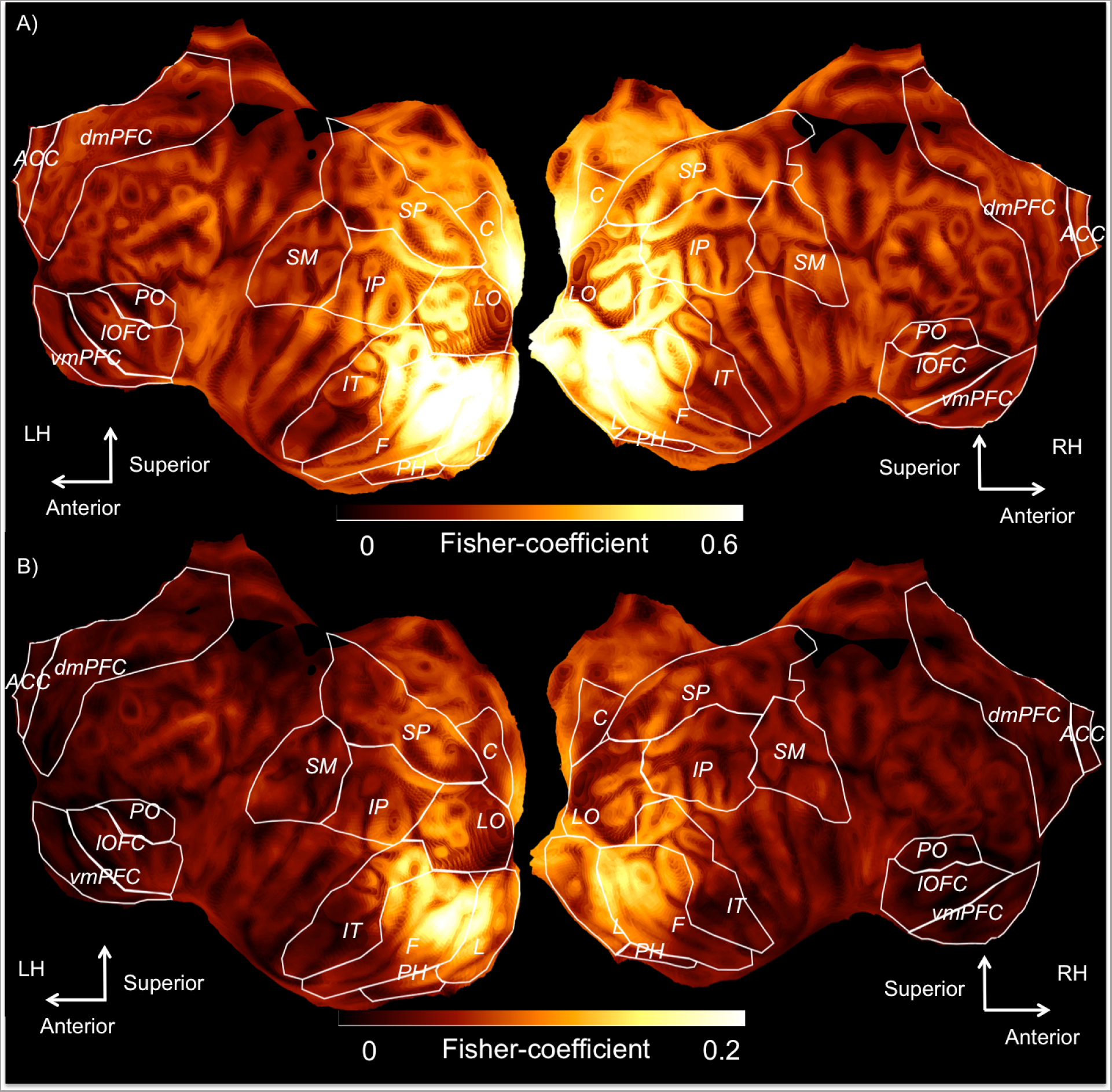
Information transmission analyses during normal conscious perception (Decoder Construction) (A) and during Neural-Reinforcement (unconscious occurrences of Target) (B). In a searchlight procedure^16^, sparse linear regression was used to predict the linearized likelihood of Target representation in ventral temporal cortex from multivoxel patterns within each sphere. Plotted in the MNI space are the mean Fisher-transformed correlation coefficients representing the accuracy of this prediction. The maximum of data scales were adjusted to reflect significant voxels determined using a permutation test. Overall transmission was lower during Neural-Reinforcement than in normal conscious viewing. ACC: anterior cingulate cortex; dmPFC: dorsomedial prefrontal; lOFC: lateral orbitofrontal; TP: temporal pole; SM: supramarginal gyrus; IT: inferior temporal; F: fusiform; PH: parahippocampal; PO: parsobitalis; SM: supramarginal; LO: lateral occipital; C: cuneus; SP: superior parietal; IP: inferior parietal.

The fact that our successful intervention could be conducted in a double-blinded fashion represents a significant methodological advancement, since conventional psychological and neurofeedback interventions can rarely be conducted with such experimental rigor. This could help decrease attrition and prevent the use of conscious “safety signals” or “safety behaviors” during exposure, which are known to interfere with the efficacy of exposure-based therapies^13^.

However, it has also been suggested that unconscious fear reduction interventions might only impact the physiological reactivity to feared stimuli, but not necessarily behavioral outcomes and subjective experiences^14^. Strictly speaking, such physiological responses may be more correctly identified as threat-related, rather than reflecting the subjective experience of fear itself^15^. While this does not undermine the validity and intended scope of the current study, it would be advantageous and interesting to test in the future how our intervention procedure may be combined with conventional (conscious) psychotherapeutic treatments to produce synergistic and long-lasting effects on clinical outcomes as well as conscious experiences.

In sum, we have exploited an opportunity to apply recent advances in fMRI decoding to move one step further towards our ultimate goal of creating a method for an unconscious brain-based psychotherapy for anxiety disorders. This study provides the first evidence that physiological fear responses to specific, subclinical, naturally-occurring, phobias can be reduced unconsciously with Hyperalignment Decoders, completely outside of the awareness of human subjects. The staggering progress of current neuroimaging decoding technology^5,17,18^, combined with in-scanner virtual reality experiments^19,20^, may mean that we can eventually extend our approach to other forms of fear, such as acrophobia, anxiety induced by public speaking, fear associated with specific persons or episodic memories, etc. In particular, for post-traumatic stress disorders (PTSD), it has been estimated that as few as 2% of patients receive sufficient treatment in the form of traditional exposure^21–23^. Our unconscious brain-based method may eventually alleviate this challenging and critical problem of patient attrition.

## Methods

The experiment was conducted in six sessions carried out on different days: Day 0: Decoder Construction session; Day 1: Pre-Reinforcement session and Neural-Reinforcement; Day 2-4: Neural-Reinforcement and Day 5: Neural-Reinforcement and Post-Reinforcement session (Fig. 3a).

### Participants

Thirty participants (8 females; Mean age = 24.0; SD = 3.97) took part in a Decoder Construction session and were included in the hyperalignment procedure. Seventeen participants (5 females; Mean age = 21.92; SD = 1.54) also took part in the Neural-Reinforcement experiment. Participants in the Neural-Reinforcement experiment first went through the Decoder Construction session and were selected for Neural-Reinforcement if they reported, on a 7-point Likert scale, “high” or “very high” fears of at least two animals included in the database. We predetermined the number of participants based on a previous study^7^. The experiment was conducted in a double-blinded procedure (i.e., neither the participants nor the experimenters were aware of the Target category of the Neural-Reinforcement procedure). Three patients (3 females; Mean age = 26.33; SD = 7.77) also took part in a Decoder Construction session. Patients were diagnosed using the Structured Clinical Interview for DSM-IV conducted by three medical doctors trained in psychiatric assessment. Diagnoses were established by inter-rater agreement. All participants provided written informed consent and the study was approved by the Institutional Review Board of Advanced Telecommunications Research Institute International (ATR), Japan.

### Decoder Construction session and hyperalignment

Each participant underwent a 1-hour fMRI Decoder Construction session, which allowed us to construct the Hyperalignment Decoders. A total of 3,600 images (90 images per category) of 30 animals categories and 10 object categories were presented for 0.98 seconds and grouped in chunks of 2, 3, 4 or 6 images of the same category (see Supplementary Methods and Extended Data Fig. 7). Trials were organized in six runs of 600 trials interleaved with short breaks. To optimize attention to each image’s category, participants were asked to perform a 1-back task, i.e. to report when the image category changed with a button press. The sequence of presentation was pseudo-randomized and fixed across participants to facilitate hyperalignment. In order to allow high-pass filtering of the fMRI data, chunks within each category were organized so that their period was smaller than 120 seconds.

We constructed the Target decoders for a designated participant from the data of 29 ‘surrogate’ participants. This method was chosen to determine how effective this procedure could be if participants were never exposed to the Target category during the Decoder Construction session, as would be ideal in a clinical setting. To do so, we iteratively performed a new hyperalignment for each category and for each participant.

To do this, we first set aside, for each designated participant, the multivoxel patterns elicited by the Target category plus an equal number (90 trials) of randomly-selected patterns associated with the remaining Non-target categories. This was done to prevent circularity^24^, as the set-aside data for the designated participant was later used to test the accuracy of the Hyperalignment Decoders. The remaining data from all participants were used to carry out hyperalignment and to develop the abstract common decoder space. This procedure involved determining a set of geometric transformations (rotation, translation and scaling) that brought data from the native space of each participant (where individual voxels are dimensions) into a common space where brain representations could be optimally aligned between participants. Importantly, this transformation can be reversed such that data from the common space can be projected back into the native space of participants. Another important feature of hyperalignment is that new data can be transformed into the common space even if they were previously withheld from hyperalignment. We capitalized on both of these features to build our training dataset; we brought all of the data from all participants (which included the Target category previously set aside) back into the native space of the designated participant via first transforming it into the common space. This allowed us to construct the Hyperalignment Decoder in the native space of the designated participant. The decoder was trained to discriminate Target from Non-target trials using the data of 29 surrogate participants and was tested on the data of the target participant (see Fig. 1).

To acquire the data necessary for this procedure, the fMRI images acquired during the Decoder Construction session were realigned to the first fMRI image, coregistered, and motion corrected in SPM 12 (Statistical Parametric Mapping; http://www.fil.ion.ucl.ac.uk/spm). The anatomical mask of the ventral temporal cortex (fusiform, lingual/parahippocampal and inferior temporal cortex) was hand-drawn using tksurfer, and using the Freesurfer (http://surfer.nmr.mgh.harvard.edu/) segmentation as a guideline. Voxels in the ventral temporal region were detrended, and then deconvolved using the least-square separate approach^25,26^.

This method allows us to iteratively construct a general linear model for each trial in the design that includes one parameter modeling the current trial, and two parameters modeling all other trials in the design. Via this method we were able to deconvolve the trials of our rapid-event related design in order to obtain parameter estimates for each individual trial, which were used in the remaining steps of the procedure. Based on previous procedures^3^, hyperalignment was conducted in pyMVPA (http://www.pymvpa.org) on a subset of 1,000 ventral temporal voxels that were the most associated with image categories; these relevant voxels were identified using an *F*-test conducted on all of the categories in the dataset but the Target category. We used the Procrustean transformation (scaling, translation, and rotation) and the common space aggregation by averaging; the hyperalignment was achieved by first iteratively projecting the individual dataset into a common space, and then iteratively projecting the original datasets on the intermediate common space processed in the first level.

After hyperalignment, data from all categories and from all participants were projected in the common space and back in the native space of the designated participant. We used sparse logistic regression^27,28^ to further select the most discriminant voxels for the Target category (on average 141.9 voxels; SEM = 4.0) and to identify a linear hyperplane that would maximally separate voxel patterns associated with the Target category from those associated with the randomly-selected Non-target images. We trained these decoders on the data from the ‘surrogate’ participants averaged within runs (6 runs) and categories. The training dataset therefore consisted of 348 multivoxel patterns distributed over the 1,000 ventral temporal voxels. The performance of the Hyperalignment Decoders was then tested by using the multivoxel patterns of the designated participant that had been held out from hyperalignment and Decoder construction (i.e., the 90 trials of the Target category and 90 trials randomly selected from the Non-target categories). Fig. 2a shows the contrast of the sparse-logistic-regression weights on each voxel between animal versus object categories (see Supplementary Methods). Fig. 2b shows the accuracies of the Hyperalignment Decoders iteratively constructed with 29 ‘surrogate’ participants and averaged over the 30 participants and the 40 object categories.

### Pre- and Post-Reinforcement sessions

To assess changes in physiological fear responses, we used brief visual presentations of animals from two feared categories (i.e., Target and Control categories), before and after the Neural-Reinforcement sessions.

Each session included the presentation of 30 images divided in 2 short blocks: 10 images of the Target condition, 10 of the Control condition, 5 images of a neutral animal (determined by a 7-point Likert scale) and 5 images of a neutral object. These presentations were carried out in the MRI scanner while electrodermal activity (skin conductance response) and fMRI images were acquired. The images presented during Pre-Reinforcement and Post-Reinforcement were never presented during the hyperalignment procedure and were created following the same procedure (see Supplementary Methods). Each trial included the presentation of a fixation cross for 3 to 7 seconds (M = 5, SD = 2), presentation of the image for 6 seconds, and then a blank screen for 4 to 12 seconds (M = 8, SD = 3). Each block started with 20 seconds of rest followed by the presentation of the image of a neutral object (e.g., a chair). The next two images were randomly set to be from the Target and Control category, and their order was counterbalanced between blocks. The remaining images were then presented randomly during the rest of the block. To estimate physiological fear responses, we built on previous methodologies^7,29^, and calculated skin conductance and amygdala responses during the first two trials of the two feared categories within each block. These mean responses were then baseline-corrected by subtracting the mean responses to the neutral animal category.

#### Skin conductance response

The skin conductance response was determined during a time window beginning 1 second and ending 5 seconds after stimulus presentation. The amplitude of the response was determined by subtracting the mean baseline activity during the 2 seconds preceding the onset of the image from the maximum value in this time window. Responses smaller than 0.02 microsiemens (μS) were recoded as 0 following previous procedures^29,30^. Responses were square root transformed to correct for the skewness of the distribution of skin conductance responses^7,31^.

We carried out a two-level mixed effect model on skin conductance response with trials nested in subjects. The model included 3 fixed effects (Time, Condition, and Time * Condition), and the coefficients of Time and Conditions were allowed to have a random component to correct for potential clustering of errors within subject^32^. Simple effect analyses were also carried by splitting the data by Condition. One participant was removed from this analysis because of a problem with the acquisition of the skin conductance data. One block of data was also removed for one participant due to a problem with image presentations.

#### Amygdala response

The amygdala ROI was determined using the structural mask resulting from Freesurfer segmentation (http://surfer.nmr.mgh.harvard.edu). The data were first realigned and coregistered in SPM. A general linear model was conducted that included, for each block, predictors (and their time derivatives) for the first 2 Target animals, the first 2 Control animals, the baseline animals, the baseline objects, and the motion parameters. The predictors were convolved with the standard canonical hemodynamic response function. Mean ROI parameter estimates were extracted for the Target and Control categories using the MarsBar toolbox^33^ implemented in SPM and baseline-corrected by subtracting the baseline condition (i.e., the neutral animal) from the mean ROI parameters.

The data were analyzed with a repeated-measures ANOVA with 2 within-subject factors of Time (Pre- and Post-Reinforcement) and Condition (Target and Control) followed by paired-sample *t*-tests (see Fig. 3b). One block was removed for one participant due to a problem with image presentations.

### Neural-Reinforcement session

The aim of the Neural-Reinforcement sessions was to allow participants to associate a reward with the activation of the neural representation of a feared animal in the ventral temporal cortex (Target category). To select participants for Neural-Reinforcement, we chose individuals who self-reported “high” or “very high” fear of at least two animals in our database. One animal category of these two was randomly selected (by a computer) to be the Target of the intervention and the other to be the Control condition. This within-subject control condition allowed for a double-blinded procedure, since neither the experimenters nor the participants were aware of the Target of the intervention during the procedure. Since participants frequently reported high fears of more than two categories (on average 4.8 feared categories; S.E.M. = 0.86), we selected, within the “high” and “very high” fear categories, the two categories presenting the Hyperalignment Decoders with the highest accuracies. For two participants, the multivoxel representations of these two categories were also correlated with one another (*r* > 0.25 both within-subject and in the hyperalignment data of surrogate participants). In these situations, we selected (1) the feared category with the highest accuracy and (2) the next feared category in terms of accuracy that was not correlated with the first category selected (*r* < 0.25).

Online Neural-Reinforcement was conducted across five sessions on five different days. Following previous procedures^7,9–12^, each trial began with an induction period (6s) during which participants were instructed to “activate a pattern in their brain” in order to maximize the size of a subsequently-presented feedback disc (i.e., the diameter of the inner grey circle) (Fig. 2c). Online decoding was achieved using the Hyperalignment Decoder for the Target category, while the decoder for the Control category was never used for reinforcement. The diameter of the circle during the Feedback period was a function of the ‘induction likelihood’ of the Target category, i.e. how likely it was that the Hyperalignment Decoder predicted the Target category from the multivoxel pattern in ventral temporal cortex. Participants were informed that their monetary gain would be a function of the overall success in correctly activating brain patterns (i.e., activation likelihood) during each session, but – critically – they were not told what the Target multivoxel pattern represented. Previous studies have shown that participants are able to learn to consistently activate the relevant voxel patterns without ever consciously understanding how to do so^7,9–12^. Induction and Feedback periods were separated by a 6s period that allowed us to account for the hemodynamic delay and to perform the online decoding on data from the time window corresponding to the Induction period (3 TRs). During each trial, the real-time data were realigned to match the coordinates of the Decoder Construction session by realigning to the first EPI image acquired during this session. Motion correction was then applied online using the Turbo BrainVoyager software (Brain Innovation, The Netherlands). Polynomial trends were removed from the activity of individual voxels and the signal was standardized voxelwise using the 20 sec waiting period at the beginning of each block. The induction likelihood of the Target condition was then computed based on the average multivoxel pattern during the 3 TRs corresponding to the Induction period, and displayed back to participants using a disc with a radius proportional to the computed likelihood. The feedback disc was presented inside a ring indicating the disc’s maximum possible size. At the end of each block, participants were informed of the monetary gain associated with their performance during the reinforcement sessions. The maximum monetary reward for each block was 300 yen (∼ USD $2.50). Participants received their earned monetary rewards at the end of each day.

Participants went through five days of Neural-Reinforcement with the Pre-Reinforcement and Post-Reinforcement sessions conducted respectively on Day 1 and Day 5. During Days 2 through 4, participants completed an average of 10.13 blocks. Day 1 and Day 5 presented fewer reinforcement blocks (M = 5.21 SD = 1.6), as Pre-Reinforcement and Post-Reinforcement sessions were also conducted on these days.

#### Information transmission analyses

The Hyperalignment Decoder likelihood computed based on voxels in the ventral temporal region could be associated with the transmission of information to other brain regions. This process could occur both during the actual presentation of the Target category during Decoder Construction as well as during pattern induction in the Neural-Reinforcement sessions. In order to compare the flow of information between the Decoder Construction phase and induction, we used an information transmission analysis^7,9,10,12^. This analysis uses a searchlight approach^16^ in which a sphere (radius = 15mm; M = 266 voxels) is iteratively centered around each voxel of the gray matter mask in the native space of each participant. Within each sphere, sparse-linear-regression machine learning classification is used to determine if it is possible to use the activity of the voxels within the sphere to predict, on a trial-by-trial basis, the linearized induction likelihood for the Target category predicted by the Hyperalignment Decoder constructed in the ventral temporal cortex. The predicted values are then correlated with the true linearized likelihood of the ventral temporal decoders. The correlation coefficients for each sphere are then Fisher-transformed and assigned to the central voxel of the sphere. The coefficients were then projected in the MNI space, and smoothed using a Gaussian filter (FWHM = 6 mm). The results of the information transmission analysis are presented on the MNI brain during Decoder Construction (Fig. 4a) and during Neural-Reinforcement (Fig. 4b). PyCortex^34^ was used for data presentation in Fig. 4.

For the information transmission analysis during Decoder Construction, the preprocessing of the data was the same as for the hyperalignment data, and the data were deconvolved using the least-square separate method. The 90 trials of the Target category were selected plus 90 other trials randomly selected from the remaining image categories (Non-target images). The linearized induction likelihood of each trial was computed using the same ventral temporal decoder used during induction. Within each sphere, the training of the sparse linear regression was cross-validated by training on 5 blocks of the Hyperalignment Decoder Construction procedure and testing on the remaining block.

Regarding the information transmission analysis during induction, we selected the same time window used to compute induction likelihoods during the Neural-Reinforcement procedure (i.e., we shifted the induction time window by 3 TRs to account for the hemodynamic response and averaged the 3 TRs that covered the induction period) and the induction likelihood was the same as the likelihood calculated and presented to the participant (via the feedback circle) during Neural-Reinforcement. Here, the sparse linear regression conducted within each sphere was trained on the decoder construction data and tested on the induction data.

To correct for multiple comparisons, a permutation test was conducted on the linearized correlation coefficients projected in the MNI space^7,35^. One thousand iterations of randomly permuted data were used to determine the distribution of the maximum t-values. Using this distribution, the critical value for significance was determined as the t-value associated with a 5% false-positive rate.

## Acknowledgements

We thank K. Nakamura and M. Miuccio for their help in scheduling and conducting the experiment, N. Hiroe and H. Moriya for assistance with equipment, Y. Shimada and A. Nishikido for operating the fMRI scanner, B. Maniscalco, M. Peters, and B. Odegaard for comments on the manuscript and M. Craske, and M. Sun, for discussions about the experiments. The study was conducted in the ImPACT Program of Council for Science, Technology and Innovation (Cabinet Office, Government of Japan). This work was partially supported by ‘Brain machine Interface Development’ under the Strategic Research Program for Brain Sciences supported by the Japan Agency for Medical Research and Development (AMED). This study was also partly funded by the US National Institute of Neurological Disorders and Stroke of the National Institutes of Health (grant no. R01NS088628 to H.L.). V.T-D. is supported by a fellowship from the Fond de Recherche du Québec - Santé (FRQS).

## Author Contributions

V.T-D., A.C., J.D.K., H.L., and M.K. designed the study, V.T-D., J.D.K., and A.C. implemented the experiment, V.T-D., A.C., and T.C. conducted the experiment, V.T-D., A.C., M.K., and H.L. analysed the results and V.T-D., A.C., T.C., J.D.K., M.K., and H.L wrote the manuscript.

---

**Extended Data Fig. 1.**
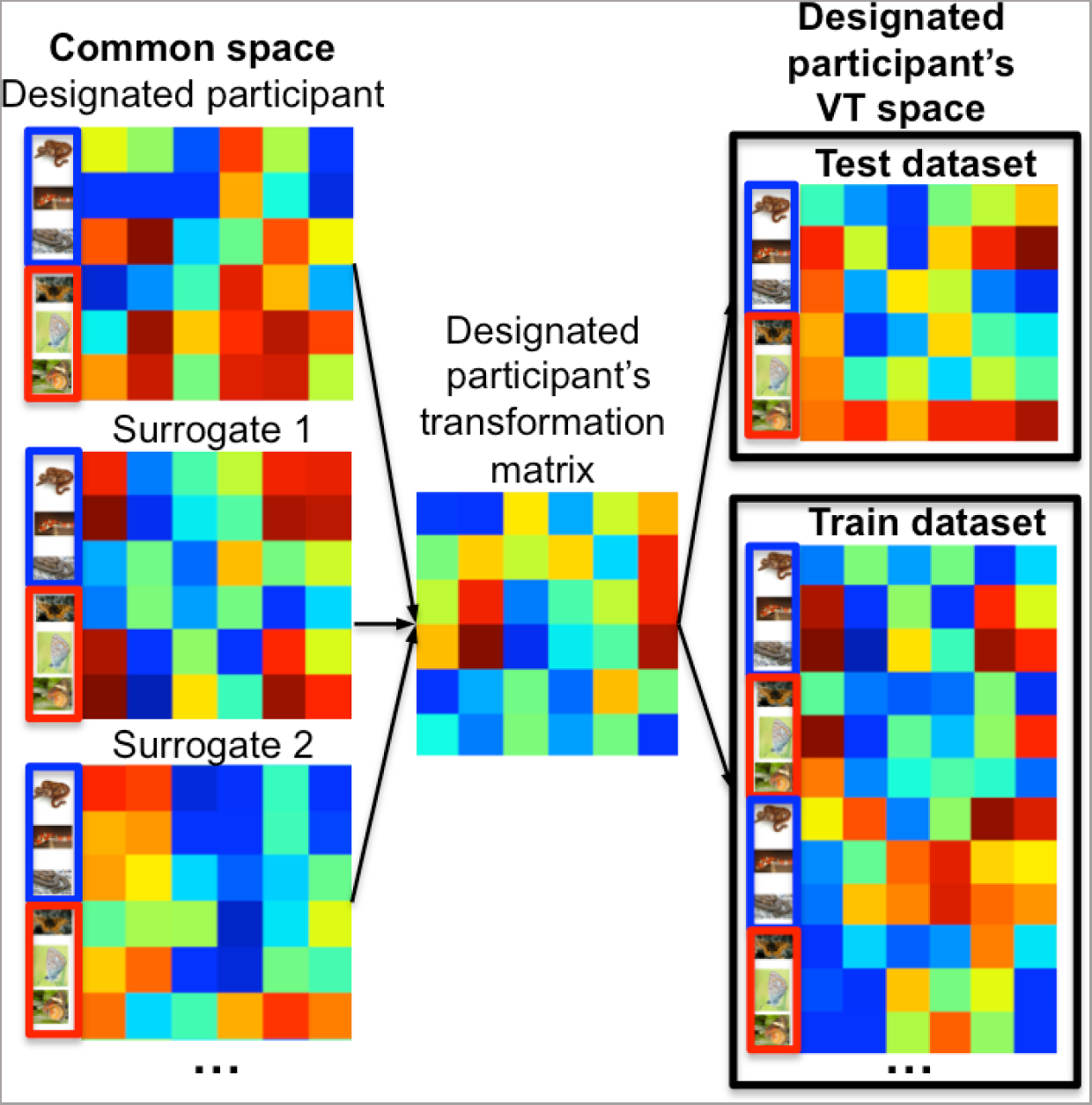
The Training and Test datasets used for Hyperalignment Decoders construction. After the hyperalignment procedure, all data for the designated participant and ‘surrogate’ participants were brought in the common space (left). By inverting the transformation matrix for the designated participant, the data of the ‘surrogate’ participants were brought into the native space of the designated participant. This approach allows us to develop a dataset for training the Hyperalignment Decoders that includes the data of Target (blue) and Non-target (red) categories from 29 surrogates. The Hyperalignment Decoder was tested on the 90 individual trials of the Target category of the designated participant as well as 90 other Non-target trials randomly selected from the remaining categories (right). The test dataset was not included in the hyperalignment or in the training of the Hyperalignment Decoder.

**Extended Data Fig. 2.**
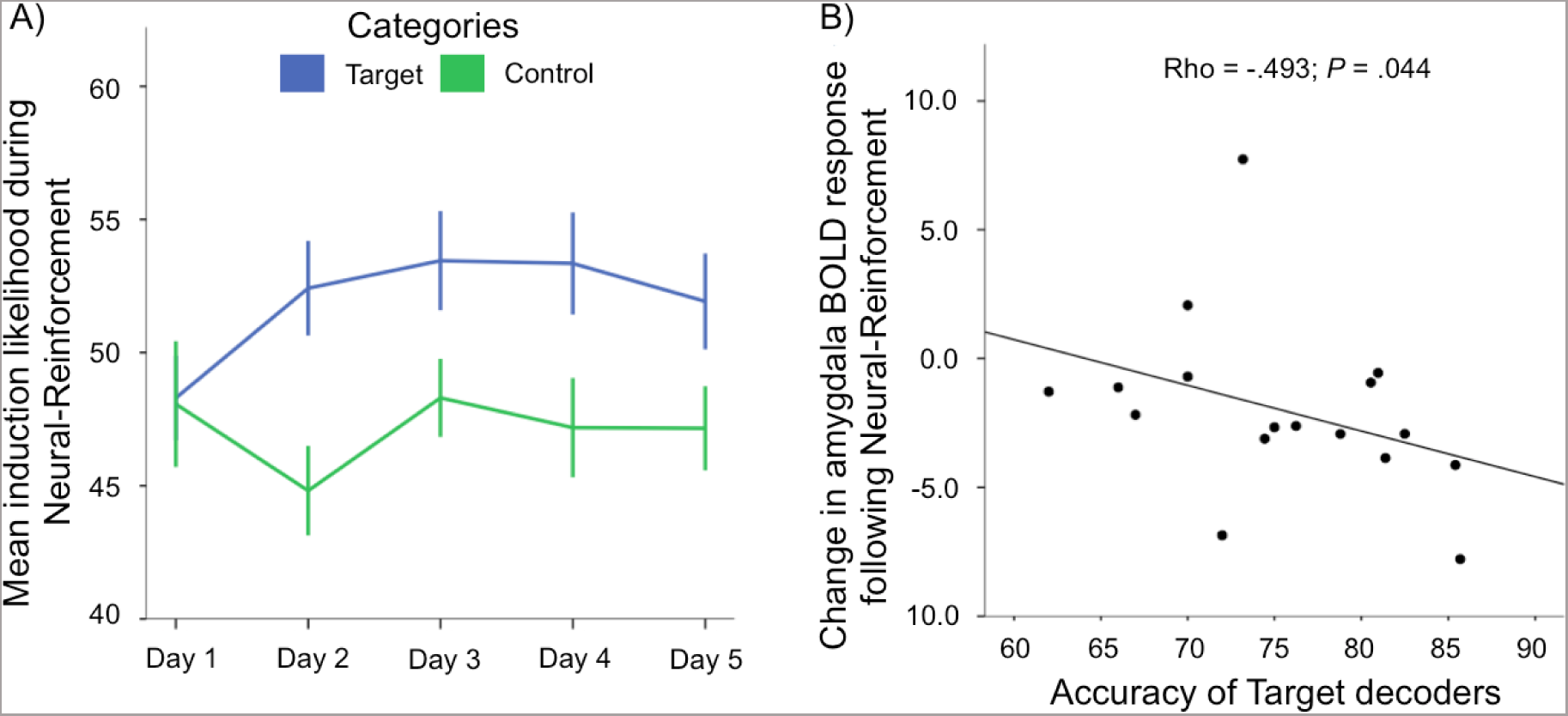
A) Mean induction likelihood of the Target and Control categories during Neural-Reinforcement. The induction likelihood of the Target category was, on average, higher than that of the Control category. Both categories presented similar likelihoods on Day 1 and significant difference on Day 2, Day 3 and Day 5. A marginally significant difference was also observed on Day 4. This indicates that a learning effect occurred during Neural-Reinforcement. (see Supplementary Discussion: Induction accuracy during Neural-Reinforcement) B) Across participants, the better the accuracy of the decoders, the more effective was the change in amygdala response in the Target condition following Neural-Reinforcement. BOLD: blood-oxygen-level dependant. Error bars are +/- 1 S.E.M. (see Supplementary Discussion: Hyperalignment Decoders used during Neural-Reinforcement).

**Extended Data Fig. 3.**
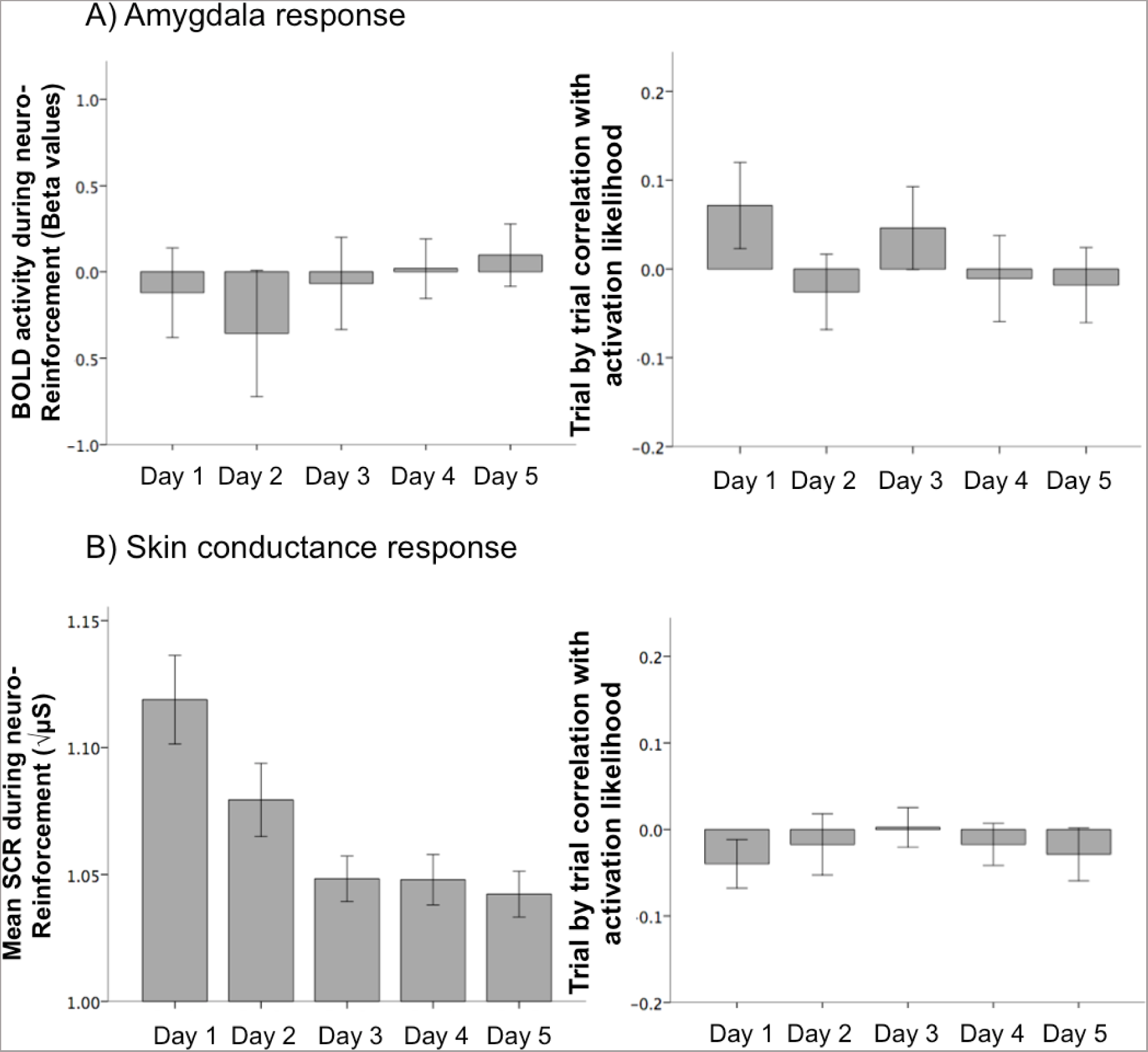
Amygdala (A) and skin conductance response (B) during Neural-Reinforcement. A) ROI analyses indicated no amygdala response during Neural-Reinforcement (left panel) and no correlation between amygdala response and the induction likelihood of the Target category during Neural-Reinforcement (Fisher-transformed spearman rho, right panel). ROI analyses were conducted using the structural masks of the amygdala (see Supplementary Discussion: Amygdala response during Neural-Reinforcement). B) Participants showed skin conductance responses during Neural-Reinforcement that decreased with time (Fisher-transformed Spearman rho, left panel). However, on a trial-by-trial basis this skin conductance response was not associated with the induction likelihood of the Target category (right panel). Error bars are +/- 1 S.E.M. (see Supplementary Discussion: Skin conductance response during Neural-Reinforcement).

**Extended Data Fig. 4.**
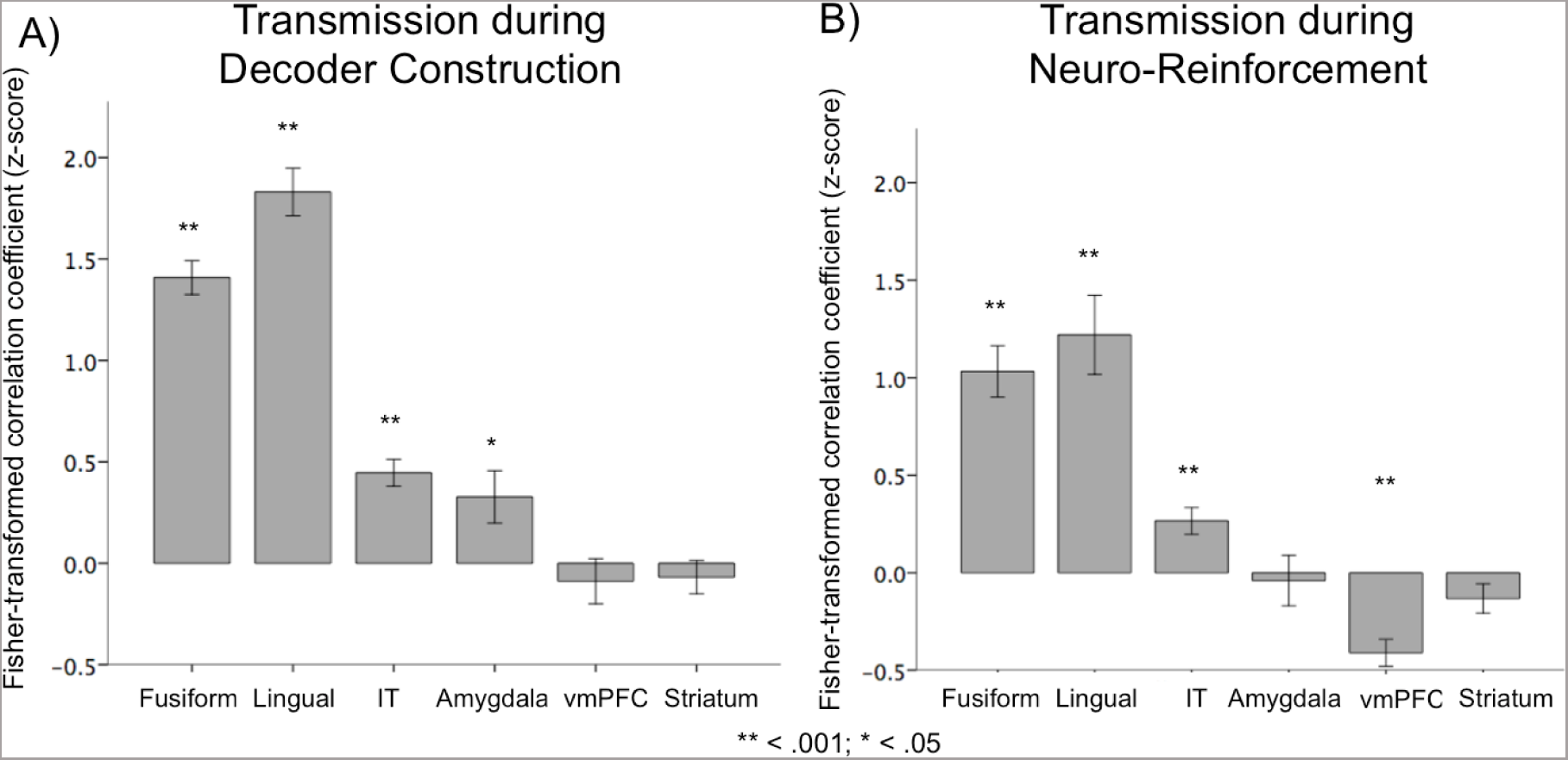
ROI analyses of the Fisher-transformed correlation coefficients of the information transmission analyses during (A) Decoder Construction and (B) Neural-Reinforcement. IT: inferior temporal; vmPFC: ventral medial prefrontal cortex. Error bars are +/- 1 S.E.M. (see Supplementary Discussion: ROI analyses of the transmission of information).

**Extended Data Fig. 5.**
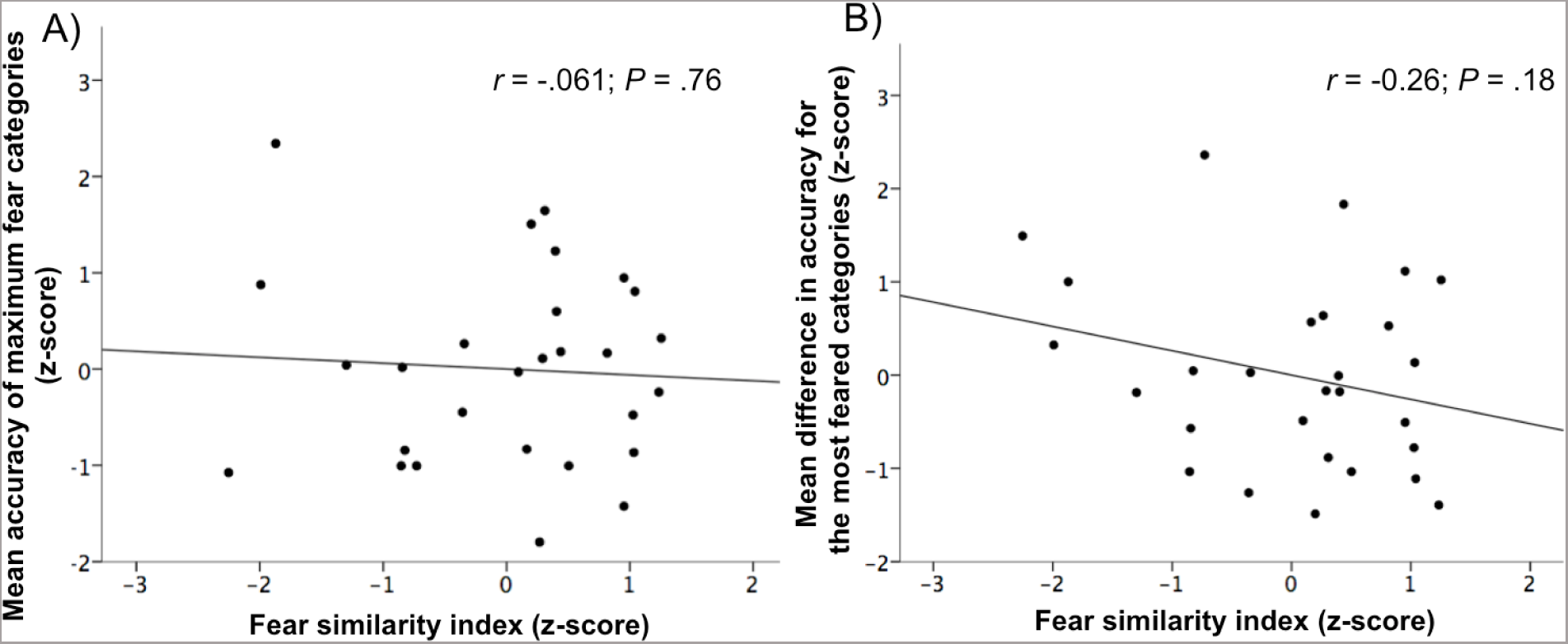
The effect of similarity between individual fear profiles and the average fear profile on Hyperalignment Decoder accuracy. The Fear Similarity Index indicates how similar the fear profile (i.e., pattern of fear ratings for various categories) for a particular participant was to the average fear profile for all other participants in the hyperalignment database. This Fear Similarity Index was computed by correlating, for each participant, fear ratings for the 30 animals with the average fear ratings for all the other participants. (A) The Fear Similarity Index was not associated with the accuracy of decoders for the most feared categories. (B) The Fear Similarity Index was also not associated with improvement in decoding accuracy afforded by the hyperalignment procedure relative to within-subject decoding accuracy. (see Supplementary Discussion: The association between fear and the accuracy of Hyperalignment Decoders).

**Extended Data Fig. 6.**
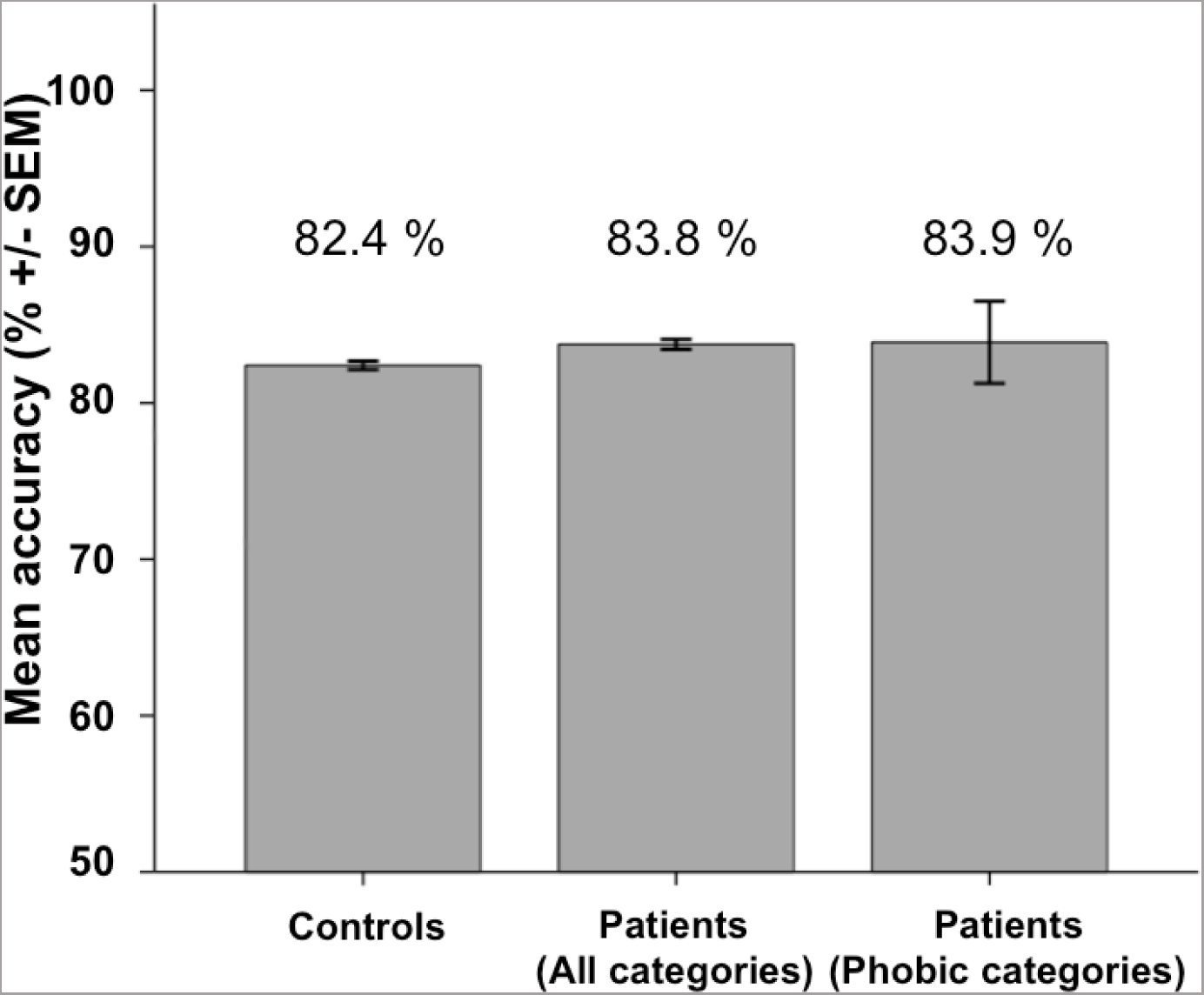
Preliminary data indicating that specific phobia diagnoses do not affect the accuracy of the Hyperalignment Decoders. As indicated in Fig. 2b, Hyperalignment Decoders present high levels of accuracy when trained and tested on control participants. Such high accuracies are also observed when the Hyperalignment Decoders are trained on control participants and tested on patients diagnosed with specific phobias (N = 3). This was observed for all categories (30 animals and 10 objects) as well as for the categories that are the object of the specific phobia (here: snake, mouse, and cockroach). Error bars are +/- 1 S.E.M. (see Supplementary Discussion: The association between fear and the accuracy of Hyperalignment Decoders).

**Extended Data Fig. 7.**
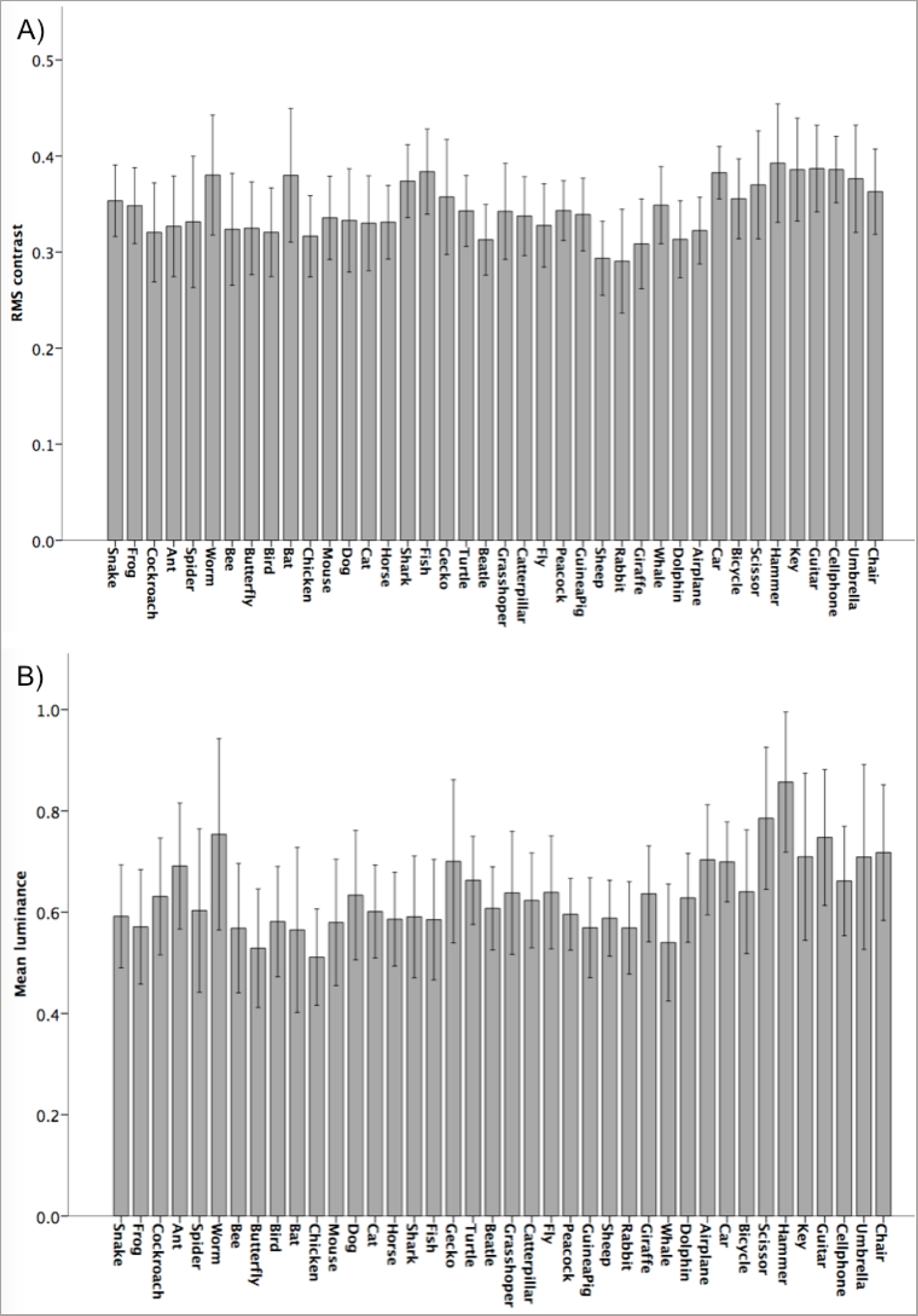
Low level visual features of stimuli presented during Decoder Construction averaged by categories. A) Mean luminance. B) mean RMS contrast. Error bars are +/- 1 SD. (see Supplementary Methods: Stimuli of animal and object categories).

